# Musical training does not enhance neural sound encoding at early stages of the auditory system: A large-scale multisite investigation

**DOI:** 10.1101/2024.09.02.610856

**Authors:** Kelly L. Whiteford, Lucas S. Baltzell, Matt Chiu, John K. Cooper, Stefanie Faucher, Pui Yii Goh, Anna Hagedorn, Vanessa C. Irsik, Audra Irvine, Sung-Joo Lim, Juraj Mesik, Bruno Mesquita, Breanna Oakes, Neha Rajappa, Elin Roverud, Amy E. Schrlau, Stephen C. Van Hedger, Hari M. Bharadwaj, Ingrid S. Johnsrude, Gerald Kidd, Anne E. Luebke, Ross K. Maddox, Elizabeth W. Marvin, Tyler K. Perrachione, Barbara G. Shinn-Cunningham, Andrew J. Oxenham

## Abstract

Musical training has been associated with enhanced neural processing of sounds, as measured via the frequency following response (FFR), implying the potential for human subcortical neural plasticity. We conducted a large-scale multi-site preregistered study (n > 260) to replicate and extend the findings underpinning this important relationship. We failed to replicate any of the major findings published previously in smaller studies. Musical training was related neither to enhanced spectral encoding strength of a speech stimulus (/da/) in babble nor to a stronger neural-stimulus correlation. Similarly, the strength of neural tracking of a speech sound with a time-varying pitch was not related to either years of musical training or age of onset of musical training. Our findings provide no evidence for plasticity of early auditory responses based on musical training and exposure.

---

Music is universal across human societies and can serve multiple functions^1^. In many Western cultures, parents may seek early musical education for their children, sometimes in the hope that the skills learned from musical training may transfer to other aspects of life. A considerable body of research has examined the potential link between such early training and enhanced perceptual and cognitive skills, with mixed results^2–7^. However, there is more consensus that musical training from an early age is associated with enhanced responses to sound at different levels of the auditory pathways, from the auditory brainstem^8–14^ to the midbrain and cortex^15–19^.

A widely reported neural signature of musical training involves the frequency following response (FFR), a scalp-recorded potential measured using electroencephalography (EEG). The FFR is thought to reflect stimulus-entrained neural responses to periodic sounds, primarily from subcortical levels of the auditory system, but also potentially reflecting some cortical contributions, especially at frequencies of 100 Hz and below^20,21^. Several studies have reported that musicians exhibit stronger FFR spectral encoding of sounds than non-musicians either for the fundamental frequency (F0) or the upper harmonics of the stimulus, and that musicians’ spectral encoding is more robust to interference from background noise^8,9,12–14,22^. Such findings have important implications: if the relationship is causal, it implies the potential for plasticity of subcortical auditory function, at least in childhood (when musical training typically begins). Even if the results from these cross-sectional studies do not stem from musical training-training induced changes in subcortical auditory responses, an alternative interpretation linking the strength of responses in the early stages of the auditory pathways (perhaps genetically influenced) to future musical success would also be highly intriguing.

Enhanced FFRs in musicians have been demonstrated in numerous studies for a variety of stimuli, including speech and non-speech sounds^8,9,12–14,22^. Two widely used types of stimuli are speech sounds with vowels that are sustained in pitch over time (e.g., /da/, Figure 1a) and speech sounds that vary in pitch over time (e.g., the Mandarin utterance, /mi3/, which dips and rises in F0 over time, Figure 1b,f, black line)^23^. The fidelity of neural encoding can be assessed either in the time domain (Figure 1c,d) or in the spectral domain (Figure 1e,f). One seminal study found that musicians had stronger spectral encoding for the upper harmonics of /da/ presented in background noise relative to non-musicians^10^.

**Figure 1.**
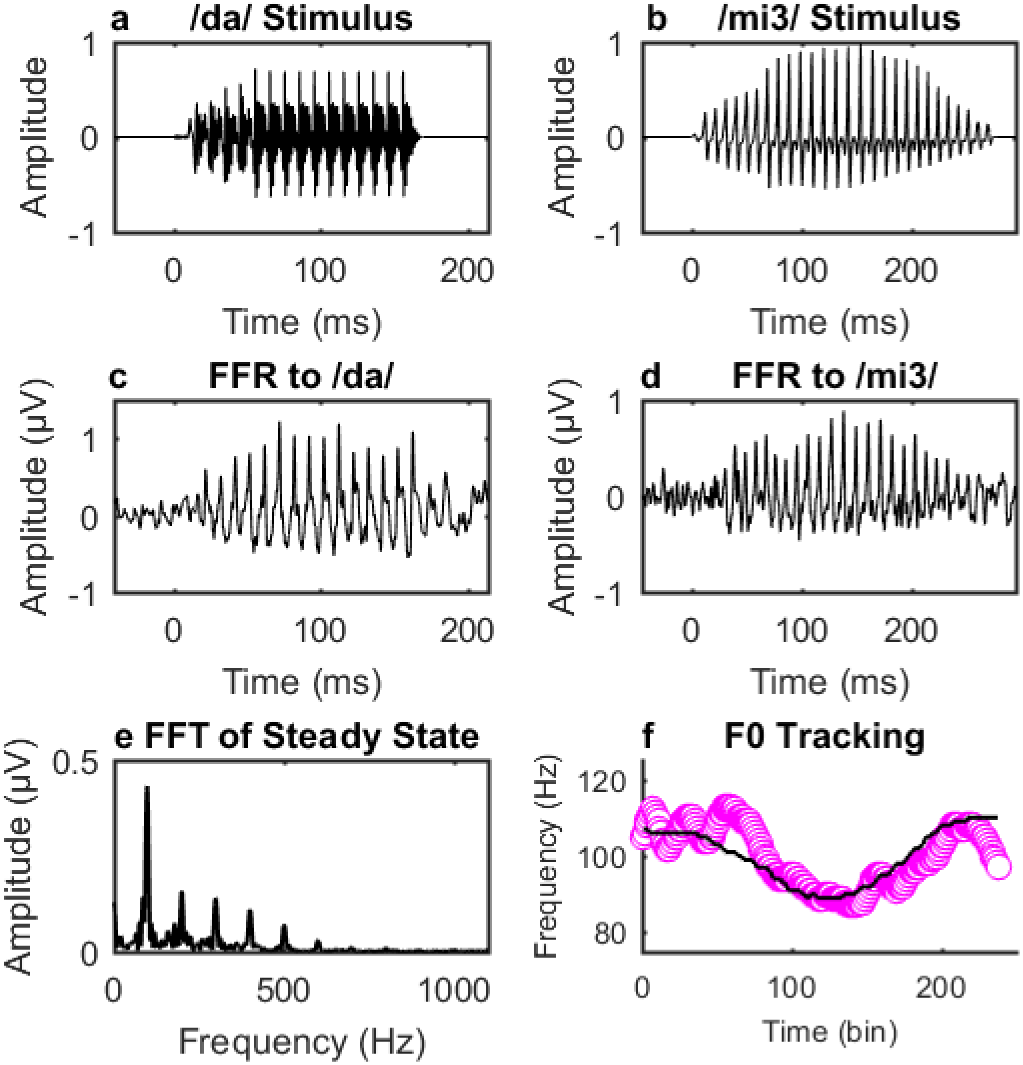
Stimuli and sample neural responses. The acoustic waveforms for /da/ (a) and /mi3/ (b) are shown, along with example FFRs in the time (c,d) and spectral domain (e,f) for one individual participant from the present study. The FFT of the steady-state portion of the FFR to /da/ (e) shows the spectral encoding for the F0 (100 Hz) and upper harmonics. Previous research found that musicians had stronger spectral encoding for the upper harmonics to /da/ presented in babble, relative to non-musicians, and stronger stimulus-to-response correlations between the vowel portion of the /da/ stimulus (50–170 ms) and the steady-state portion of the neural response (60–180 ms)^10^. The F0 of the /mi3/ stimulus (f, black line) varies in F0 over time (pink: neural F0 tracking). The original study found that musicians (relative to non-musicians) had stronger stimulus-to-response correlations between the F0-trajectory of the stimulus and the neural response and that F0-tracking fidelity was related to age of onset of musical training.

Furthermore, the correlation between the averaged time waveform of the neural response and the stimulus waveform was greater in musicians, indicating that musicians’ FFRs were more robust to the presence of background noise. Another influential study showed that native English speaking musicians had a stronger representation of the time-varying F0 for a Mandarin word presented in quiet^12^, as quantified by the correlation between the F0 contour of the stimulus and neural response (Figure 2f). Importantly, the strength of the neural representations of the stimulus F0 was negatively related to the age at which musicians began their musical training. This relationship suggests that earlier onset of musical training produces stronger neural responses. Together, these two studies have had a major impact, as evidenced by the high number of citations (a combined total of 1770, as of the time of writing), and form the early foundation of the subsequent body of positive evidence for enhanced early neural encoding of sound in musicians^24^.

**Figure 2.**
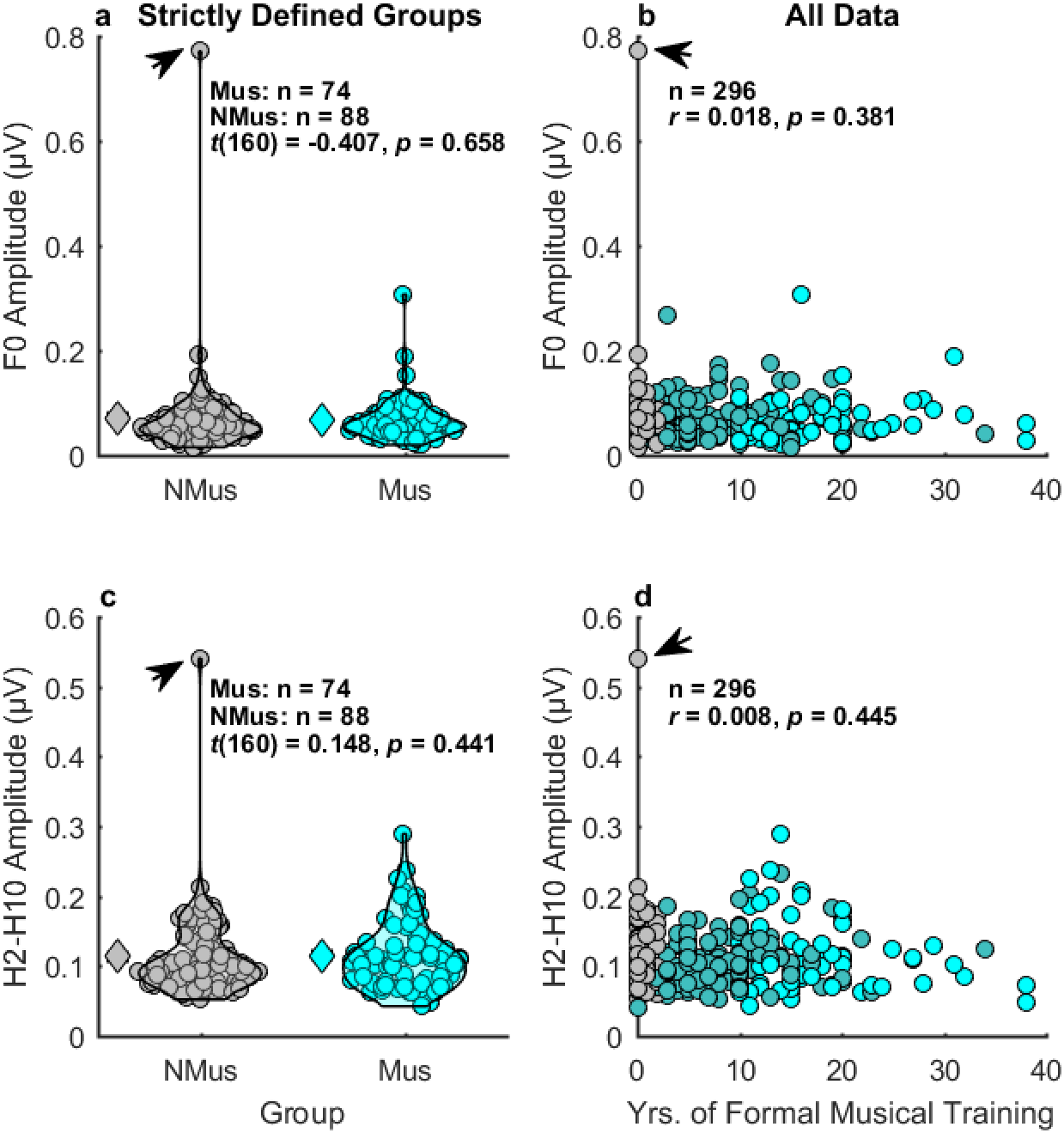
Musical training is not related to enhanced spectral encoding of the F0 (a,b) or upper harmonics (c,d) of /da/ syllable in background noise. No effect was found when comparing the strictly defined groups of musicians and non-musicians (a,c), and no relationship was observed in the broader sample between neural encoding strength and years of musical training (b,d). The null effects of musical training remained after removing one outlier non-musician with unusually strong spectral encoding (indicated by arrow; see Supplementary Figure 1 for plots with outlier removed). Black outlines: 1D kernel density estimates (KDEs); Diamonds: Average data; Circles: individual data; NMus: non-musicians (grey); Mus: musicians (cyan). Participants in neither of the strictly defined groups are shown in dark cyan.

Although current evidence supports the notion that musicians exhibit advantages in the neural encoding of sound^24^, several factors complicate the interpretation of this advantage. First, most reports have been based on relatively small samples of listeners^10,12,25^ with dichotomous samples that often represent extreme ends of the musical spectrum (i.e., untrained people compared to musicians with many years of experience). Limiting the sample in this manner is beneficial from an experimenter standpoint, in that it increases the likelihood that between-group differences will be detected, but such disparate groups may differ in many other ways besides musicianship, such as socioeconomic status or personality^2,26^, limiting the generalizability of the findings^27^. Moreover, most studies have been conducted on young (college-aged) adults, although there are a growing number of studies examining aging effects^28–33^. Another complicating factor is that the definition of the terms “musician” and “non-musician” has varied between studies, leading to the possibility that any differences in outcomes between studies may reflect, in part, differences in years and nature of training, age training began, and degree to which musical training or activity is maintained.

In the present study, we attempted to replicate the two seminal findings of a musician advantage for spectral neural encoding of speech sounds, described above^10,12^, across a large sample of participants at six different sites. In addition to replicating, we also extended these studies in several important ways: (1) All sites conducted both studies, allowing for a high-powered aggregate sample. (2) All participants at each site took part in both studies, allowing for the strength of neural encoding between studies to be compared within the same participants. (3) Both age and musical training varied continuously, increasing the generalizability of the findings relative to previous studies, while still allowing for a direct replication by maintaining a subset of participants who fit the most stringent criteria for the definitions of musicians and non-musicians. (4) The methods and primary analyses were preregistered before beginning data collection^34^, limiting researcher degrees of freedom when analyzing the data. (5) Additional data were collected on factors that have been found to co-vary with musical training, including personality (i.e., the openness to experience factor of the ‘Big 5’ personality scale) and socio-economic status^26^, which could be used in exploratory analyses or by other researchers in future analyses. The results provide important insights into the replicability and robustness of the musician advantage in early neural processing of sound across the adult lifespan.

## Results

### Spectral encoding of the syllable /da/ in multi-talker babble

#### Group comparisons

The first study we sought to replicate^10^ compared the neural spectral encoding of the vowel portion of the syllable /da/ (Figure 1a,c,e), embedded in multi-talker babble, in 16 musicians and 15 non-musicians, as they passively watched a silent video. In that study, musicians were found to have enhanced encoding of the upper harmonics (H2-H10) but not the F0. We first conducted a direct replication in a subset of our participants that represented the extreme ends of the musical training spectrum (74 musicians and 88 non-musicians). As shown in Figure 2 (a,c), there was no significant difference in spectral encoding as indexed by FFRs between musicians and non-musicians for either the F0 [*t*(160) = -.407, *p* = .658, *d* = -.064] or upper harmonics [*t*(160) = .148, *p* = .441, *d* = .023] of the same /da/ syllable used by the original study. The lack of effect persisted after removing one non-musician outlier who had unusually strong spectral encoding [Supplementary Figure 1; F0: *t*(159) = .684, *p* = .248, *d* = .108; Upper harmonics: *t*(159) = .909, *p* = .182, *d* = .144]. Bayesian analyses indicated the data were 7.81 and 5.24 times more likely to occur under the null hypothesis than the alternative hypothesis, for encoding the F0 (BF_+0_ = .128, % error < .0001) and upper harmonics (BF_+0_ = .191, error = ∼.057), respectively. Excluding the outlier non-musician also provided moderate support that the data originated under the null (F0: BF_+0_ = .315, % error < .0001; upper harmonics: BF_+0_ = .404, % error < .0001). The results remained robust across a wide range of widths of the Bayesian prior (Supplementary Figure 2). These results show no relationship between musical training and the FFR for /da/ in background multi-talker babble for neither the F0 nor the upper harmonics.

The spectral amplitudes of the FFR decrease with age^35^, which may have added unexplained variance to our data, potentially obscuring a musician effect. Across the full sample of participants, we confirmed that age was related to poorer spectral encoding of both the F0 and upper harmonics of /da/ in babble (F0: *r* = -.246, *p* < .0001, one-tailed; upper harmonics: *r* = -.173, *p* = .001, one-tailed; Supplementary Figure 3). Our subsamples of highly experienced musicians and inexperienced non-musicians did not differ significantly in age [*t*(160) = −1.3, *p* = .196; outlier excluded: *t*(159) = −1.36, *p* = .176; two-tailed tests]. Exploratory ANCOVAs also confirmed that musicianship was not related to enhanced spectral encoding for either the F0 [*F*(1,159) = .444, *p* = .506, *ƞ* ^2^ = .003] or upper harmonics [*F*(1,159) = .004, *p* = .947, *ƞ* ^2^ < .0001] even after adjusting for effects of age. As before, these findings remained the same when excluding the outlier non-musician with the strongest spectral encoding [F0: *F*(1,158) = .123, *p* = .726, *ƞ_p_*^2^ = .0008; upper harmonics: *F*(1,158) = .494, *p* = .483, *ƞ_p_*^2^ = .003].

The original study^10^ also reported that musicians had significantly greater stimulus-to-response correlations than non-musicians, suggesting that musicians’ neural encoding of speech sounds was more robust to noise than that of non-musicians. We attempted to replicate this finding by conducting an independent-samples *t*-test on the z-transformed stimulus-to-response correlations between musicians and non-musicians. We found no musician advantage using our pre-planned analyses (Fig. 3a), with the average trend in the opposite direction to that predicted [*t*(160) = -.716, *p* = .763, *d* = -.113]. The Bayes factor was BF_+0_ = .106 (error = ∼.002), meaning the data are 9.43 times more likely to come from the null than the alternative hypothesis, providing moderately strong support for the null (Supplementary Figure 4). Our analysis followed the traditional approach of selecting the time-lag between stimulus and response that produced the greatest correlation, whereas the original study limited the stimulus lag times to the range of 6.9–10.9 ms prior to adjusting for site-specific delays between the onset of the trigger and the arrival time of the stimulus at the ear canal. An exploratory analysis using the original study’s approach also showed no musician advantage [*t*(160) = .074, *p* = .471, *d* = .012] with moderate support for the null hypothesis (BF_+0_ = .18, error ∼.04%).

**Figure 3.**
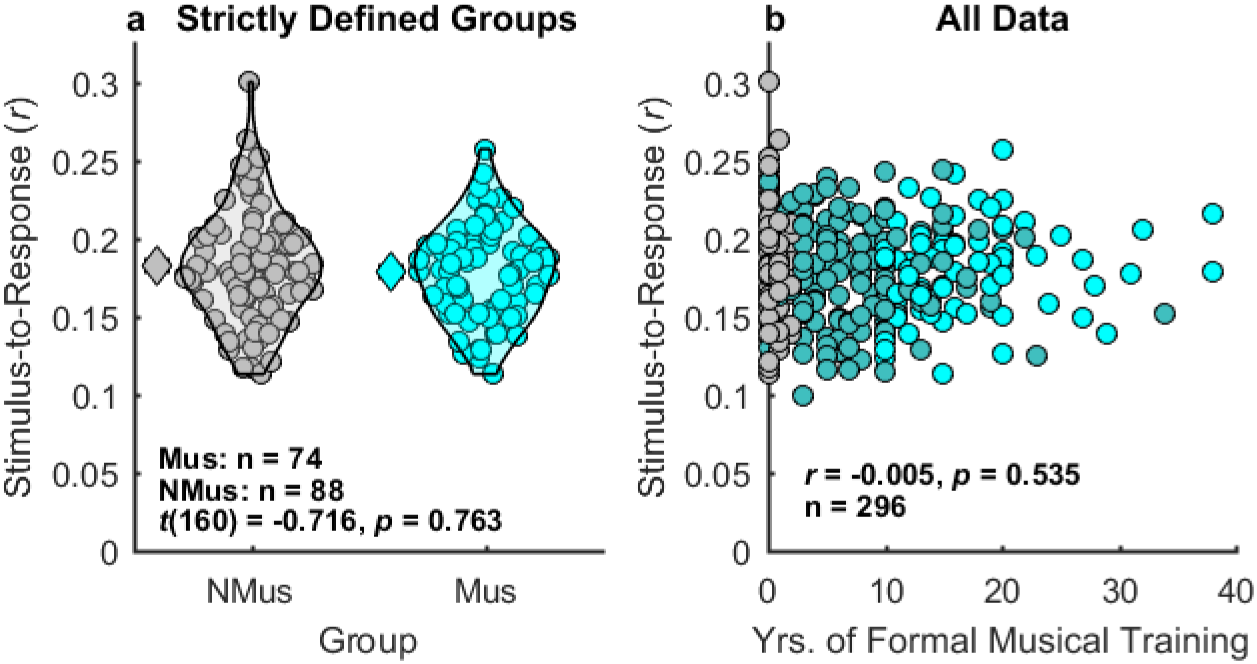
Musical training is not related to more robust encoding of speech in background sounds. The untransformed stimulus-to-response correlations are plotted for visualization purposes; the z-transformed data were used in analyses. Black outlines: 1D KDEs; Diamonds: Average data; Circles: individual data; NMus: non-musician (grey); Mus: musician (cyan). Participants in neither of the strictly defined groups are shown in dark cyan.

#### Continuous comparisons

Next, we tested whether years of formal musical training, as measured across the entire cohort of participants, was correlated with enhanced neural encoding of /da/ in babble, quantified as: (1) the strength of F0 encoding (Figure 2b), (2) the strength of encoding the upper harmonics (Figure 2d), and (3) the transformed stimulus-to-response correlations (Figure 3b). None of these preregistered hypotheses were tested in the original study^10^, but they are extensions of the underlying hypothesis that musical training is associated with enhanced neural representation of speech in noisy backgrounds. A Bonferroni-corrected criterion for significance (α = .017) was preregistered. Years of formal musical training was not related to enhanced spectral encoding for the F0 (*r* = .018, *p* = .381; non-musician outlier excluded: *r* = .094, *p* = .054, Supplementary Figure 2) or upper harmonics (*r* = .008, *p* = .445; non-musician outlier excluded: *r* = .048, *p* = .204); similarly, years of musical training did not offset the masking effects of background noise on speech encoding, as quantified via the stimulus-to-response correlation (*r* = -.005, *p* = .535). An exploratory analysis limiting the cross-correlations to the site-specific adjusted lag windows of 6.9-10.9 ms, as in the original study, also showed no relationship between the adjusted stimulus-to-response correlation and years of formal musical training (*r* = .029, *p* = .312). Further exploratory partial correlations controlling for age confirmed no significant effects of years of musical training (see Supplementary Information).

### Neural encoding of linguistic pitch contours

#### Group comparisons

The original study^12^ measured EEG responses while participants listened passively to the Mandarin word /mi3/ as they watched a silent video. The stimulus varied in F0 over time between 89 to 110 Hz (Figure 1f). The original study found that the F0 stimulus-to-response correlation (i.e., the Pearson correlation between the F0 contour of the stimulus and the neural response) was significantly greater in 10 musicians than in 10 non-musicians. We attempted to replicate this finding on the subset of our participants that met the strict criteria of musician and non-musician (68 musicians and 77 non-musicians; Figure 4a) and found no musician advantage [*t*(143) = 1.32, *p* = .094, *d* = .22], with the data about 1.4 times more likely to occur under the null than the alternative hypothesis (BF_+0_ = .715, error < .0001%, Supplementary Figure 5). The effect remained non-significant after excluding a musician-group outlier with the poorest neural encoding [*t*(142) = 1.63, *p* = .052, *d* = .273] but resulted in a Bayes factor very close to 1 (BF_+0_ = 1.14, error < .0001%), implying no strong evidence for either the alternative or the null hypothesis.

**Figure 4.**
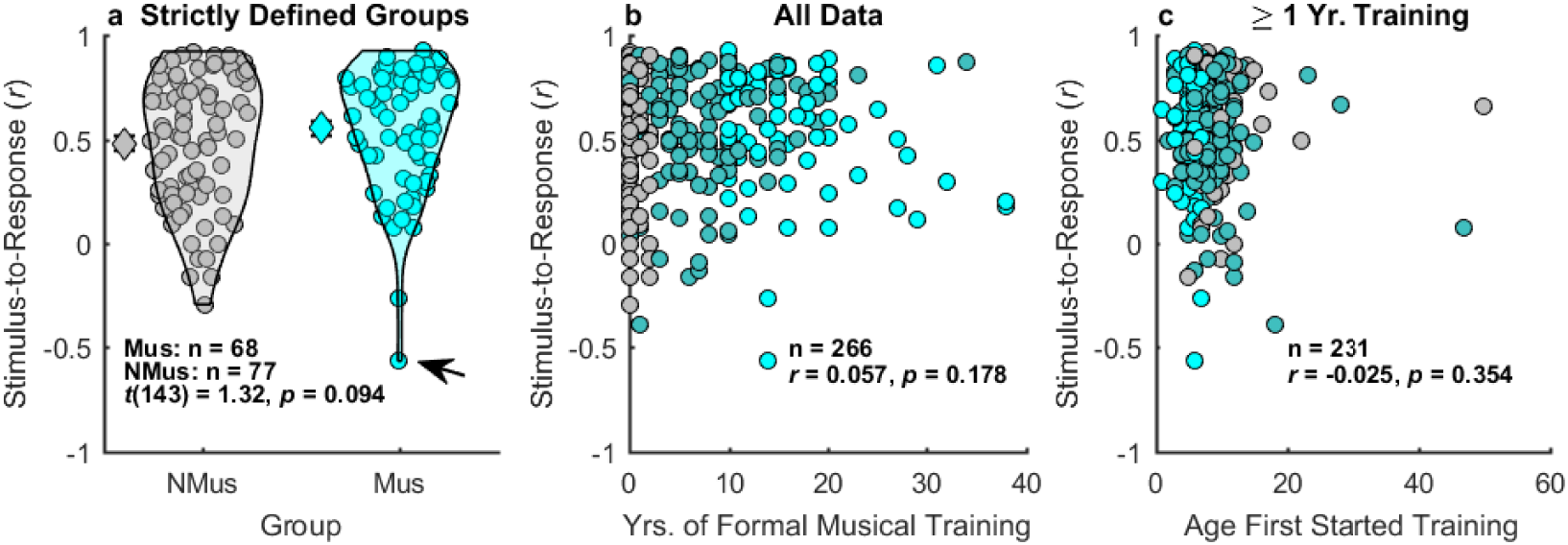
Neural encoding for time-varying F0 is unrelated to duration and age of onset of musical training. No significant difference was observed between the strictly defined groups of musicians and non-musicians (a). There was also no relationship between neural tracking of the F0 and years of formal musical training (b) or age of onset of musical training (c). Diamonds: Average data; Circles: individual data; NMus: non-musician (grey); Mus: musician (cyan). Participants in neither of the strictly defined groups are shown in dark cyan; Arrow indicates outlier data point.

To test if age was masking any effect of musicianship on F0-tracking fidelity, an exploratory analysis tested for a musician advantage while adjusting for age. There was still no benefit of musicianship on F0-tracking, either with [*F*(1,142) = 1.6, *p* = .208, *ƞ* ^2^ = .011] or without [F(1,141) = 2.47, *p* = .118, *ƞ* ^2^ = .017] the musician outlier with the poorest encoding.

#### Continuous comparisons

The original study^12^ found that the age of onset of musical training as well as years of musical training were correlated with the fidelity of F0 tracking, assessed via the F0 stimulus-to-response correlation. We used the full cohort of participants (excluding those with 0 years of musical training) to test whether age of onset of training was related to F0 tracking (Figure 4c). We also tested the hypothesis that F0 tracking improves with years of musical training by calculating the correlation between years of formal musical training and the F0 stimulus-to-response correlation (including those with no musical training, as in the original study^12^; Figure 4b). The preregistered criterion for significance included Bonferroni correction for two comparisons (α = .025). Unlike the original study, which included 16 participants, we found no relationship between the age of onset of musical training and the fidelity of neural encoding of time-varying stimulus F0 (Figure 4c; *r* = -.025, *p* = .354), with the data 8.77 times more likely to occur under the null than the alternative hypothesis (BF_-0_ = .114; Supplementary Figure 6). Furthermore, the relationship between years of formal musical training and the fidelity of F0 tracking was not significant (Figure 4b; *r* = .057, *p* = .178), with moderate evidence that the data originate under the null hypothesis (BF_+0_ = .192). Exploratory partial correlations controlling for age (Bonferroni-corrected α = .025) confirmed no relationship between age of onset of musical training and F0 stimulus-to-response correlations (*r_p_* = -.005, *p* = .469) or years of formal musical training and F0 stimulus-to-response correlations (*r_p_*= .057, *p* = .177). However, F0-tracking did worsen with age (*r* = -.173, *p* = .002), consistent with the expected degradation of spectral neural encoding with age^35^.

### Comparing neural responses between measures

Most participants (n=263; Mus = 68; NMus = 74) completed both the /da/ test and the /mi3/ test and met all inclusion criteria for analyses (see Methods). We compared the strength of neural encoding between tests to examine whether neural tracking for the F0 of speech in quiet is related to FFRs for encoding for speech in babble (Supplementary Figures 7,8). Exploratory analyses demonstrated that the stimulus-to-response correlation for encoding the F0 of /mi3/ in quiet was weakly related to F0 spectral encoding for /da/ in babble (*r* = .133, *p* = .016, Bonferroni-corrected α = .0125), but this effect did not reach significance and was driven by one outlier non-musician with very strong encoding for both measures (outlier removed: *r* = .072, *p* = .122). There was no association between neural encoding for /mi3/ in quiet and the upper harmonics of /da/ in babble (*r* = .071, *p* = .125; outlier removed: *r* = .009, *p* = .442), and there was still no association once one additional outlier musician, with poor F0 tracking but strong encoding for the upper harmonics, was removed (*r* = .068, *p* = .138). While measures between studies were unrelated to one another—potentially because dynamic F0 tracking is somewhat different from overall strength in spectral encoding, or because the stimuli were different—we did find a relationship between strength of spectral encoding for the F0 versus upper harmonics within the same stimulus (/da/ in babble: *r* = .585, *p* < .0001; outlier non-musician removed: *r* = .3, *p* < .0001).

### Does musical ability account for failures to replicate?

Our criteria for the groups of “musician” and “non-musician” were at least as strict as in both original studies, so the failure to replicate any musician advantages in neural encoding is unlikely to be due to our group definitions. However, musical expertise or aptitude, rather than years of training, may be a more sensitive measure for detecting differences in neural encoding of sound^36^. We tested this directly by correlating an objective measure of musical ability (same/different melody discrimination)^37^ with our four measures of neural encoding fidelity: (1) spectral encoding for the F0 and (2) upper harmonics for /da/ in babble, (3) stimulus-to-response correlations for /da/, and (4) F0-tracking for /mi3/ in quiet. There was no relationship between the objective measure of musical ability and any of the four measures of sound neural encoding (Figure 5; see Supplementary Figure 9 for results with the non-musician outlier excluded).

**Figure 5.**
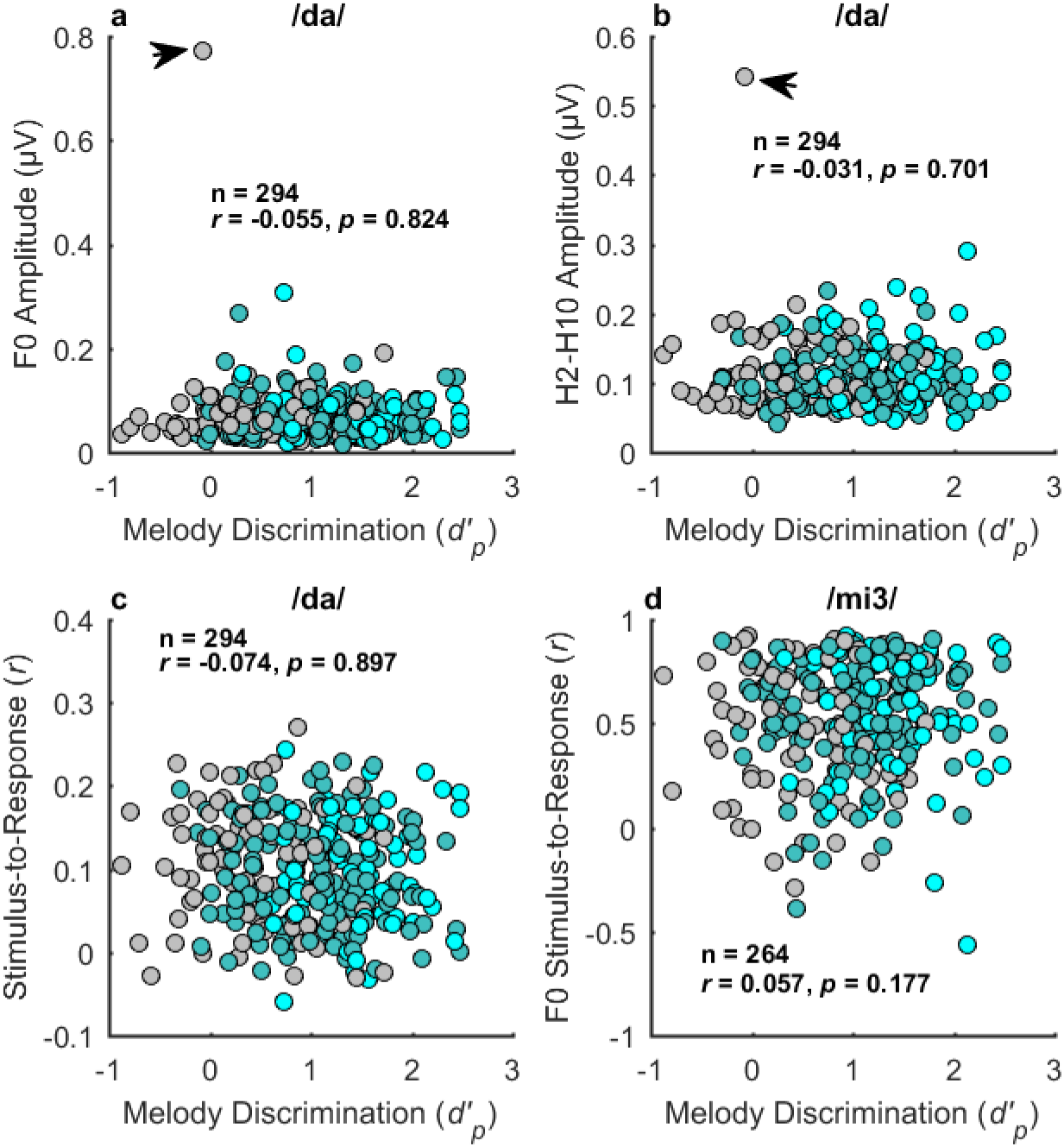
Musical ability is unrelated to sound neural encoding for /da/ in babble (a-c) or /mi3/ in quiet (d). Musical ability was assessed using an objective measure of melody discrimination^37^ and quantified using a non-parametric estimate of sensitivity, *d’_p_*^38,39^, where higher *d’_p_* indicates better performance. Grey circles: non-musicians; Cyan circles: musicians; Dark cyan circles: Participants in neither of the strictly defined groups; Arrow points to outlier data.

## Discussion

Our large-scale replication and extension tested the robustness and generalizability of the widely accepted finding that musicians have enhanced neural encoding of sound, as assessed with the FFR – an electrophysiological index of the fidelity of neural encoding in the early stages of the auditory pathways^24^. The results were consistent across all five direct-replication analyses (Table 1): There were no significant musician advantages, despite using the same stimuli, methods, and analyses as in the original studies. Musicians did not exhibit stronger sound spectral encoding for the upper harmonics of speech in background sounds, nor did they demonstrate enhanced tracking for dynamic changes in linguistic F0 for speech sounds in quiet. Neural encoding in musicians was also not more robust to the effects of background noise than that of non-musicians. An extension of the analyses to include years of formal musical training as a continuous variable, which increased the statistical power and the generalizability of the results, also revealed no relationship between musical training and any of our preregistered measures of neural encoding of sound.

**Table 1.** Key findings selected for direct replication, along with the effect sizes from present study. The number of citations listed for each paper corresponds to Google Scholar on July 16^th^, 2024. All five direct-replication analyses failed to replicate. Parentheses: original study sample size.

| Reported Musician Advantage | Publication | Sample Size |  | # of Citations | Original Effect Size | Current Study Effect Size |
| --- | --- | --- | --- | --- | --- | --- |
|  |  | Mus. | Non-Mus. |  |  |  |
| Enhanced spectral encoding of upper harmonics to /da/ in babble. | Parbery-Clark et al. (2009)<sup>10</sup> | n=74 (16) | n=88 (15) | 492 | $d = 1.04$ | $d = .023$ |
| Less degradative effects of babble on encoding speech (/da/). | Parbery-Clark et al. (2009)<sup>10</sup> | n=74 (16) | n=88 (15) | 492 | $d = 1.05$ | $d = .012$ |
| Enhanced F0 encoding for linguistic pitch in quiet. | Wong et al. (2007)<sup>12</sup> | n=68 (10) | n=77 (10) | 1278 | $d = 1.11$ | $d = .22$ |
| Years of musical training related to higher fidelity of neural encoding for dynamic changes in F0. | Wong et al. (2007)<sup>12</sup> | n=266 (20) | | 1278 | $r = .456$ | $r = .05$ |
| Starting a musical instrument at an earlier age related to enhanced F0 tracking. | Wong et al. (2007)<sup>12</sup> | n=231 (16) | | 1278 | $r = -.502$ | $r = -.025$ |

*Note.* Cohen's $d$ was estimated directly from the original studies using the reported groups means and standard deviations, if available; otherwise, $d$ was calculated from the reported $t$ -value and degrees of freedom.

Exploratory analyses controlling for potential effects of age also confirmed no significant effects of musical training on neural encoding of sound. This finding is especially important, as musical training has been proposed to potentially counteract the age-related declines of the fidelity of neural encoding of sound^4,28^, and even speech perception in background noise^32^. Our results confirmed that spectral encoding tends to degrade with age, but this effect appeared to be larger and more consistent for the encoding of the F0 compared to the upper harmonics. Our findings that poorer spectral encoding with age was unrelated to musical training provide no support for efforts to enhance the FFR in older listeners through musical training interventions^4,40^.

### Why did the effect of musicianship fail to replicate?

A recent review of perceptual and neural associations with musical training noted that conclusions were mixed with respect to behavioral outcomes but that all studies examining neural advantages in musicians had reported at least one significant effect^24^. Given the seeming robustness of the results, it may be surprising that our large-scale study failed to replicate any of the original findings. This apparent discrepancy may have several possible underlying causes. First, the positive results have come from a relatively limited number of laboratories, and even then the musician advantage has not been consistent among studies. For example, one study^11^, using the same /da/ stimulus as in the present study, found a musician encoding advantage in quiet, but only for the F0 and not the upper harmonics, whereas another study using the same stimulus^33^ reported a musician advantage for the upper harmonics, but not the F0, both with and without background noise. Second, the sample sizes from past studies in this field of research have all been relatively small (n < 30), raising the likelihood of false positives^41,42^. Finally, and perhaps most importantly, there is no single agreed-upon analytical technique for examining the fidelity of sound encoding using EEG. Researchers thus have many degrees of freedom related to the number of ways they can test their hypotheses^43^, and not all of the analyses may be reported in the final publication. Testing the same hypothesis many different ways increases the chance of a false positive (Type I error), highlighting the importance of preregistration of hypotheses, methods, and planned analyses prior to data collection.

An alternative reason for the failure to replicate is that the present study may have one or more false negatives (Type II errors), despite the large n and corresponding sensitivity to detect small effects. In fact, all five direct-replication analyses demonstrate small but non-significant effects in the expected direction. Based on the very small effect sizes reported in Table 1, if a musician advantage is present in the full population of musicians and non-musicians, then the effects seem likely to be too small to influence perception and behavior on an individual level. While traditional frequentist statistics can tell us whether we fail to reject the null hypothesis, Bayes Factor (BF) analyses can assess evidence for the null. Our BF analyses generally provided moderate evidence that the data originated under the null hypothesis for all direct-replication analyses, with the exception of the test comparing F0-tracking in musicians versus nonmusicians. But even here, there was no clear evidence for either the alternative or the null hypothesis.

Lastly, it is possible that we selected the wrong musician advantages to test. For example, a number of studies have claimed that musicians exhibit stronger and shorter-latency responses relative to non-musicians^8–10,28,33^. The majority of studies examining response latencies used an expert peak-picker to manually identify the peaks in the early neural responses. It would be impossible to directly replicate such analyses, as different findings between studies could be related to differences in the expertise or strategy of the peak picker(s), making a null finding difficult to interpret. Future analyses of our publicly available data could involve efforts to automate the process of peak-picking^44^ or formally assess the reproducibility of outcomes across different peak pickers^45^.

### Broader Implications

Our study is the first large-scale replication and extension of the widely accepted finding that musical training is related to enhanced neural encoding of sound. Our results do not support this claim. While the neural locus of the FFR is debated, and may contain contributions from cortical sources^15,20,21,46^, the evidence suggests that the dominant sources are subcortical for frequencies greater than 100 Hz, and originate primarily in the inferior colliculus^47–49^. One important direct implication is that phase-locked subcortical neural responses to sound are likely not nearly as plastic as previously thought, even following many years of intensive musical training starting at an early age. It may be that cortical structure and function is more susceptible to music-related interventions, although most studies so far have been cross-sectional, making it difficult to determine whether any differences are due to musical training^2^.

### Conclusions

Using sample sizes more than four times the size of the original studies, with preregistered methods and analyses, and data collected in six laboratories, we showed that the widely accepted finding that musicians have enhanced subcortical responses to sound failed to replicate. In an extension of the original studies, we also found no relationship between the fidelity of neural encoding of sound and years of formal musical training. Further exploratory analyses showed that musical training did not offset the age-related deterioration in the spectral encoding of sound. In all, none of the replication, extended, or exploratory analyses we conducted provided support for a relationship between musical training and sound neural encoding via the FFR.

Musical training is not related to an enhancement in early neural encoding of sound. Nevertheless, there are of course many important reasons why learning and playing music remains a valuable endeavor, including social connection, emotional regulation, or simply the enjoyment of music for its own sake^50^.

## Methods

### Recruitment and eligibility

Only participants who completed the full online screening, did not report encountering any audio issues, passed the auditory attention check, and indicated they would like to be contacted to participate in future lab studies were eligible for the laboratory portion of the study. Occasionally a potential participant met the recruitment criteria except that they reported audio issues, in which case, the researcher could invite them to redo the melody portion of the online screening in the lab. Online participants who reported a history of hearing loss (unless they were age 40 or older, in which case some high-frequency hearing loss was allowed; see Table 2), neurological conditions, proficiency in a tonal language (such as Mandarin or Cantonese), or who were not native speakers of North American English (i.e., did not live from birth through age 5 in a household where North American English was the primary spoken language) were ineligible for the laboratory portion of the study.

**Table 2.** Audiometric threshold criteria. All participants that completed the lab-portion of the study met the audiometric threshold criteria specified for their age group in each ear.

| Age Group | .125 - 1 kHz | 2 kHz | 4 kHz | 6 kHz | 8 kHz |
| --- | --- | --- | --- | --- | --- |
| 18-39 | ≤ 20 dB HL | ≤ 20 dB HL | ≤ 20 dB HL | ≤ 20 dB HL | ≤ 20 dB HL |
| 40-49 | ≤ 20 dB HL | ≤ 20 dB HL | ≤ 20 dB HL | ≤ 30 dB HL | ≤ 30 dB HL |
| 50-59 | ≤ 20 dB HL | ≤ 25 dB HL | ≤ 30 dB HL | ≤ 40 dB HL | ≤ 40 dB HL |
| 60-69 | ≤ 20 dB HL | ≤ 30 dB HL | ≤ 40 dB HL | - | - |

To ensure an adequate representation across age ranges, participants for the full lab study were recruited at each site to be roughly evenly distributed in each decade of age (20s – 60s, with ages 18 and 19 grouped in the 20s decade), based on the age reported in the online screening. Each site aimed to recruit 60 participants, with at least 25% of participants with no more than 2 years of any musical training and no ongoing music performance activities (i.e., they reported that they did not currently play a musical instrument, including voice), and at least 25% of participants who started playing their first musical instrument or voice by the age of 7, had completed at least 10 years of formal musical training, and reported that they currently played a musical instrument. Formal musical training was defined as group or private lessons, excluding standard elementary school activities. The remaining participants could have varying amounts of formal musical training. In this way, we could assess the effect of number of years of formal musical training as a continuous variable, while still being able to perform dichotomous comparisons (musician vs. non-musician) with at least half of our overall sample. Sites aimed to have roughly even and uniform distribution of ages and gender between these two groups, as with all other participants.

### Participants

All participants in the present study also took part in a number of behavioral tests in the lab (not reported here). Participants recruited for the in-person portion of the study underwent a pure-tone audiometric screening at octave frequencies between 125 and 8000 Hz, as well as at 6000 Hz. Because age and high-frequency hearing loss co-vary^51,52^, the maximum allowable hearing loss was titrated per decade, so that participants under the age of 40 were required to have thresholds ≤ 20 dB hearing level (HL) across all tested frequencies, but older adults could have more high-frequency loss (specified in Table 2). All participants were required to meet the audiometric criteria in both ears to participate. A total of 296 participants (115 male; 177 female; 4 non-binary), including 74 musicians and 88 non-musicians (as defined in “Recruitment and Eligibility” section), completed the syllable-in-noise study, and 295 completed the linguistic pitch study. Thirty of the participants for the linguistic pitch study did not meet the pre-specified criterion for analyses (i.e., at least one binned FFT analysis was in the noise floor, as defined by Wong et al., 2007^12^) or were unable to return to the lab to redo the study, so their data was excluded from this task, leaving a total of 265 participants (106 male; 155 female; 4 non-binary). Participant age ranged from 18 to 69 years for both studies. Most participants completed both studies; reasons for missing data are described in each site’s corresponding EEG log (https://osf.io/duq34/). The total number of participants per site for each measure, including their musical status, are provided in Supplementary Table 1.

All participants provided written informed consent and were given monetary compensation or course credit for their in-person participation. All study protocols were approved by the Institutional Review Board at the corresponding university site prior to any data collection: Boston University (4942E), Carnegie Mellon University (STUDY2018_00000367), Purdue University (1609018209), University of Minnesota (0605S85872 and 1306S37081), University of Rochester (STUDY00004020), and by the Nonmedical Research Ethics Board of the University of Western Ontario (NMREB 112604).

### Online screening

Before completing the laboratory portion of the study, all participants remotely completed an initial online screening. The purpose of the screening was to aid in recruitment of qualified participants for the lab-based portion of the study (e.g., based on age, years of formal musical training, etc.), acquire an objective measure of melody perception abilities, and obtain survey information on factors that may co-vary with musicianship (e.g., personality) for use in possible exploratory analyses.

The online screening was administered through Qualtrics, with each site completing recruitment and online data collection under the purview of their own IRB. All screening participants provided informed consent online. Participants were not compensated for participation in the screening, but they had the option to enroll in a drawing for a chance to win a gift card as an incentive for participating. All personal identifiers were removed from online data before sharing between sites, so that only the subject ID number linked the online to the laboratory data. The online measures are described below in the order that they appeared.

#### Age

Participants were asked to select their age from a dropdown menu. A reported age of younger than 18 or older than 89 led to termination of the screening.

#### Level adjustment

To help ensure that the stimuli for the online listening tasks were audible but not too loud, participants were presented with noise and instructed to adjust their volume so that it was at an audible but comfortable level. The stimulus was white noise, bandpass filtered between 200 and 1000 Hz, so that the frequency spectrum was comparable to that used in the melody task.

#### Attention check

This task helped exclude participants who were not attending or who did not have properly functioning audio on their device. To pass the attention check, participants were required to answer at least 3 of 4 trials correctly. Each trial consisted of a short sequence of 1-kHz pure tones. Each tone within a sequence was 400 ms in duration with 50-ms raised-cosine onset and offset ramps, and each tone was separated by 500 ms of silence. Participants were instructed that each trial contains between 0-9 tones, and their task was to report the number of tones they heard by selecting the corresponding number from a dropdown menu. Because there were 10 options for each trial, the probability of passing the screening by chance was very low (.0037). To minimize the duration of the task, each trial only had 1, 2, 3, or 4 tones in a sequence, with each tone-sequence option presented once. Trials had a fixed duration of 5 s, so that the entire task could be completed in less than half a minute. No feedback was provided.

#### Melody discrimination

Stimuli were from the Melody subtest of the Full Profile of Music Perception Skills (PROMs), with methods as described in Law and Zentner (2012)^37^. During each trial, participants first heard a reference melody twice in a row, followed by a comparison melody. The task was to determine whether the comparison melody was the same as or different from the reference melody, with participants selecting their answer from five possible options: “Definitely Same”, “Probably Same”, “I Don’t Know”, “Probably Different”, or “Definitely Different”. One practice trial was provided, followed by 18 data trials. Participants did not receive feedback, but they did receive their total composite score at the end of the task. The composite score provided to participants was calculated using weighted responses as described in Law and Zentner (2012)^37^, with confident correct responses (“Definitely Same” or “Definitely Different”) receiving 1 point, less-confident correct responses (“Probably Same” or “Probably Different”) receiving .5 point, and incorrect responses or “I Don’t Know” receiving 0 points. For analyses, melody discrimination performance was calculated using a bias-free estimate of sensitivity, *d’_p_*, as recommended by Strauss et al. (2023)^38^ and Whiteford et al. (2023)^39^.

#### Survey questions

A number of survey questions assessed factors related to demographics, musical engagement, socio-economic status, and hearing status. Self-report of any audio issues during the melody task was also gathered. The full set of questions is available at https://osf.io/duq34/.

#### Big Five Personality Inventory

The 44-item Big Five Personality Inventory was administered to assess personality^53,54^. This was a self-report questionnaire, where each item is rated on a 5-point response scale.

### Stimuli and procedures

Each study was designed with the purpose of measuring the same effect *in principle* as the original study. In some instances, small methodological deviations were employed for practical reasons, including to decrease the total time of the study or to increase the feasibility of the measure to be collected consistently at multiple sites.

All sites ran the tests in the same order as described below. Tests that were skipped or needed to be rerun due to technical or researcher error were noted in a study log and, whenever possible, the participant returned to complete tests with missing data.

#### EEG: Syllable in noise

To assess the fidelity of neural encoding for speech in noise, we measured EEG responses to the speech syllable /da/ (Figure 1, top row) in multi-talker babble. The /da/ had a 100-Hz F0 with 170-ms duration, presented at 80 dB SPL, as used by Parbery-Clark et al.^10^. Both the /da/ and multi-talker babble were generously shared by the principal investigator of the original study. The multi-talker babble had a 37.41-s duration (not 45 s, as mistakenly reported in the original study^10^) and looped continuously throughout the task at 10 dB below the level of the speech syllable. Participants listened passively to /da/ in multi-talker babble over insert earphones (see Table 3) while watching a silent video in a sound-attenuating booth. The /da/ was presented at alternating polarities over two blocks of 3000 trials each (6000 trials total), with an ISI of about 83 ms, so that each block lasted approximately 13 min. Participants were allowed short breaks between blocks and instructed to remain still during stimulus presentation. Data were acquired with the electrode systems and sampling rates listed in Table 3 with earlobe references. Sites with BioSemi systems ensured that the magnitude of the offset voltages were < ±30 mV before beginning data collection.

**Table 3.**
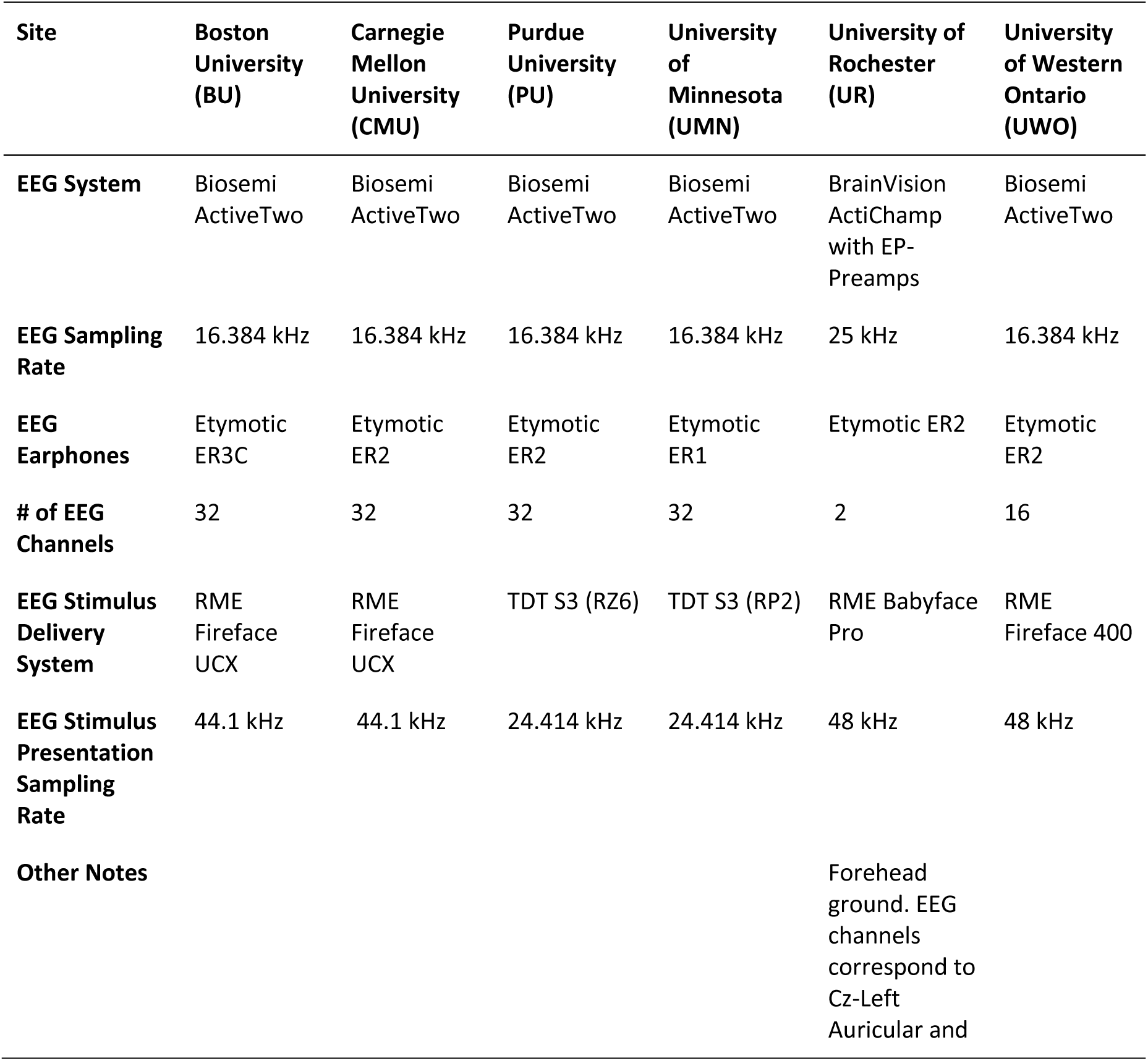
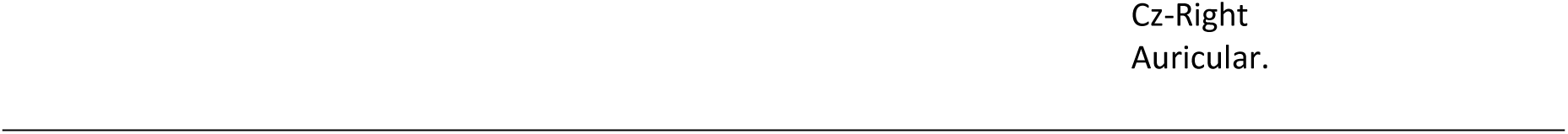
Equipment used at each data-collection site.

| Site | Boston University (BU) | Carnegie Mellon University (CMU) | Purdue University (PU) | University of Minnesota (UMN) | University of Rochester (UR) | University of Western Ontario (UWO) |
| --- | --- | --- | --- | --- | --- | --- |
| **EEG System** | Biosemi ActiveTwo | Biosemi ActiveTwo | Biosemi ActiveTwo | Biosemi ActiveTwo | BrainVision ActiChamp with EP-Preamps | Biosemi ActiveTwo |
| **EEG Sampling Rate** | 16.384 kHz | 16.384 kHz | 16.384 kHz | 16.384 kHz | 25 kHz | 16.384 kHz |
| **EEG Earphones** | Etymotic ER3C | Etymotic ER2 | Etymotic ER2 | Etymotic ER1 | Etymotic ER2 | Etymotic ER2 |
| **# of EEG Channels** | 32 | 32 | 32 | 32 | 2 | 16 |
| **EEG Stimulus Delivery System** | RME Fireface UCX | RME Fireface UCX | TDT S3 (RZ6) | TDT S3 (RP2) | RME Babyface Pro | RME Fireface 400 |
| **EEG Stimulus Presentation Sampling Rate** | 44.1 kHz | 44.1 kHz | 24.414 kHz | 24.414 kHz | 48 kHz | 48 kHz |
| **Other Notes** |  |  |  |  | Forehead ground. EEG channels correspond to Cz-Left Auricular and |  |

#### EEG: Linguistic pitch

The Mandarin word /mi3/ (the 3 denoting a dipping tone; Figure 1b,f), which means “rice,” was presented bilaterally at 70 dB SPL. Methods were adapted from Wong et al.^12^, and the stimulus was generously provided by the original authors. The word was originally recorded by a native Mandarin speaker and then adjusted in Praat^55^ to have a duration of 278.5 ms and an F0 contour ranging from 89-110 Hz. Participants passively listened to 4800 repetitions of /mi3/, divided into two blocks of 2400 trials and presented at alternating polarities (2400 trials per polarity). The ISI was about 83 ms, so that the entire task had a duration of about 30 min (15 min per block). The same electrode system, sampling rate, and active and reference electrodes was used as in the syllable-in-noise task.

### Hardware, software, and materials

With the exception of the online screening, all auditory stimuli were controlled via Matlab (R2016b). Code for tests and analyses is available on GitHub and linked to the project website on the Open Science Framework (OSF; https://osf.io/duq34/). The stimuli for the EEG studies may be available upon reasonable request by contacting the authors of the original studies^10,12^. The principal investigator from Law and Zentner (2012)^37^ should be contacted for requests to use the melody discrimination stimuli. Table 3 shows the equipment used at each test site.

### Sharing of Data

Each site was responsible for quality checking their data before sharing with the first author to ensure it was formatted in a manner consistent with the other sites. Each site was also responsible for maintaining a detailed log of each EEG session and providing the code they use to clean and format the raw data.

### EEG Data Analyses: Syllable in Noise

All single-channel preprocessing and analyses were the same as reported in Parbery-Clark et al.^10^, unless otherwise stated. The recordings from one site (UR) were down-sampled so that EEG data from all sites had the same sampling rate; this was not done in the original study but was necessary due to equipment differences. All recordings were bandpass filtered between 70-2000 Hz (12 dB/octave with zero-phase shift) and then epoched from −40 – 213 ms, where 0 ms corresponds to the stimulus onset. Trials with activity ≥ ±35 µV were treated as artifacts and removed from analyses. Epochs were baseline-corrected based on the mean potential in the pre-stimulus period. The average response was taken across the trials at each polarity, and this average was summed across polarities to minimize the stimulus artifact and the cochlear microphonic^23,56^. Only the spectral analyses were selected for replication with the aim of including fewer hypothesis tests to simplify the analysis plan.

#### Spectral encoding

The musician advantage for enhanced spectral encoding to the vowel portion of /da/ in babble was assessed using the same, fast Fourier transform (FFT) analysis methods as the original study^10^. An FFT of the steady-state portion of the EEG response (60-180 ms) was calculated for each subject, with zero padding added. The strength of harmonic encoding for the first 10 harmonics (with the first harmonic corresponding to the F0) was estimated by calculating the average spectral amplitude within 60-Hz-wide frequency bins that were centered around each harmonic. To estimate the overall strength of encoding of the upper harmonics for each subject, the average spectral amplitudes for harmonics 2-10 were summed.

#### Stimulus-to-response correlations

The preregistered stimulus-to-response correlation analysis assessed cross-correlations at all possible lag times (slightly different the original study^10^, which limited the stimulus lag to 8-12 ms) between the vowel portion of the /da/ stimulus (50 – 170 ms) and the steady-state portion of the neural response (60-180 ms), defined in the same manner as used in the FFT analysis. The maximum correlation across all lag times is referred to as the stimulus-to-response correlation, with stimulus-to-response correlations closer to 0 indicating poorer neural representations of /da/. Because correlations do not adhere to assumptions of normality, they were transformed using Fisher’s *r*-to-*z* transformation before conducting analyses; this transformation was not used by the original study^10^.

An exploratory analysis was also conducted to more precisely match the lag times tested in Parbery-Clark et al. (2009)^10^ and to account for any fixed delay between the onset of the trigger and the arrival time of the stimulus at the ear canal, which varied between sites due to equipment differences. This included accounting for any fixed delay between the onset of the trigger and the onset of the stimulus (e.g., due digital to analog conversion time) as well as the time it takes the stimulus to travel the length of the earphone tubes. The stimulus lag used in the cross-correlation analysis by the original study was 8-12 ms, which included a 1.1 ms fixed delay. We therefore limited the lag time to 6.9-10.9 ms for all sites, and then added each individual site’s fixed delay time to quantify the site-specific lag window. The cross-correlation was conducted on the neural response to the vowel (50-200 ms) and the zero-padded vowel portion of the stimulus within each site-specific lag window. The maximum correlation within the site-specific lag window for each subject is referred to as the adjusted stimulus-to-response correlation.

### EEG Data Analysis: Linguistic Pitch

Recordings were bandpass filtered between 80-1000 Hz (12 dB/octave using zero-phase shift) and then epoched from −45 – 295 ms, where 0 ms corresponds to the stimulus onset. All other data preprocessing were as described for the syllable in noise task. The F0-tracking analysis used by Wong et al.^12^ was chosen for replication because it demonstrated the most consistent evidence of a musician advantage. Preprocessing and analyses were consistent with the original study unless otherwise stated.

#### Calculating F0 tracking

The strength of F0 tracking was estimated by performing a sliding FFT analysis on the EEG response for each subject over the entire FFR period, after accounting for any fixed delay between the onset of the trigger and the arrival time of the stimulus at the ear canal. This included accounting for any fixed delay between the onset of the trigger and the onset of the stimulus (e.g., due digital to analog conversion time) as well as the time it takes the stimulus to travel the length of the earphone tubes. The time-averaged EEG response was segmented into 40-ms bins, with each bin spaced 1-ms apart. 238 bins in total were used, and a Hanning window was applied to each bin. To estimate the spectral content of each bin, an FFT on the windowed bin was conducted, with zero-padding out to 1 s. The F0 of each bin was defined as the frequency with the greatest spectral magnitude within ± .5 octave of the mean stimulus F0 (100 Hz). The latter criterion was not described in the original study, but we decided to include it based on correspondence with the first two authors of the original study for advice on how to ensure F0 tracking corresponds to the F0 rather than the upper harmonics. Any frequencies with spectral magnitudes that were not above the noise floor were excluded as possible F0s. The noise floor was calculated by performing a Hanning-windowed FFT on the average pre-stimulus period, when no stimulus was present. This method for estimating the noise floor was not described in Wong et al.^12^ but is described in Skoe and Kraus (2010)^23^ and is believed to be the method used by the original study.

The fidelity of F0 tracking was measured by comparing the F0-tracking of the EEG response to the estimated F0 of the stimulus. The /mi3/ stimulus was down-sampled to 16.384 kHz, and the same sliding FFT analysis was performed on the stimulus to assess the degree to which the FFR response matched the stimulus F0, with the first bin in the analysis beginning at time 0. The F0 within each bin was defined as the frequency with the peak magnitude within ±0.5 octaves around the true approximate stimulus F0 (100 Hz); defining such a frequency window was necessary (but not described in the original study), as the second harmonic has a greater spectral magnitude than the F0 for a majority of the stimulus. The degree to which F0 neural encoding matches the stimulus F0, or the F0 stimulus-to-response correlation, was calculated using Pearson’s *r* for each subject. Because correlation coefficients do not follow a normal distribution, each *r* was transformed to z using Fisher’s *r-*to-*z* transformation before conducting the statistical tests described below, consistent with the original study.

### Statistical Analyses

To ensure high-powered analyses for all tests, the data were aggregated across sites. All group comparisons used the same, relatively strict definition of musician and non-musician, as described in the “Recruitment and eligibility” section. This ensured that the definition of musician was generally as strict, if not more so, as the definition used in the original studies. Outliers were identified by visual inspection, and analyses were conducted both with and without outliers. Direct replication analyses used the same statistical tests as the original study. Analyses that treated years of formal musical training as a continuous variable were conducted across the full cohort of participants unless otherwise stated. As the overarching finding across the original studies was that musicianship provides an advantage to sound processing, all corresponding significance tests were one-tailed with α=.05 unless otherwise noted. Exploratory analyses that used age as a covariate were conducted using Analysis of Covariance (ANCOVA) for between-group comparisons and partial correlations for continuous comparisons.

ANCOVA statistical assumptions of linearity and homogeneity of regression slopes were tested by visually inspecting scatterplots including the regression lines between the covariate and dependent variables for each group. Homogeneity of regression slopes was also tested by ensuring the interaction between the group and co-variate had a *p*-value > .05. Homogeneity of variances was tested using Levene’s test of equality of error variances. Data were analyzed in Matlab 2016b and JASP^57^.

Exploratory Bayes factor (BF) hypothesis tests supplement the direct replication analyses in order to assess the support for the alternative hypothesis (i.e., musicians are better than non-musicians) versus the null hypothesis using the reporting standards outlined in van Doorn et al. (2021)^58^. Between-groups comparisons were assessed using Bayesian independent-samples *t* tests with a truncated Cauchy prior distribution 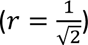 so that only positive effect sizes were examined. Bayesian Pearson correlations used a uniform prior. Robustness was assessed across a wide range of prior widths, with results reported in the supplementary materials. BF supplementary figures were created using JASP version 0.18.3 and compiled using Adobe Illustrator CS6.

#### Spectral encoding for /da/: Group comparisons

The overall strength of spectral encoding in musicians and non-musicians was assessed using two independent samples t-tests, one for the F0 and one for the upper harmonics. This analysis is identical to the original study but deviates slightly from our preregistered plan (see Supplementary Information).

#### F0 tracking for /mi3/: Group comparisons

The hypothesis that musicians would have better F0-tracking than non-musicians was tested using an independent-samples t-test on the z-transformed F0 stimulus-to-response correlations between the two groups.

#### Musical ability and sound neural encoding fidelity

Melody discrimination performance was scored by calculating *d_p_*, a non-parametric estimate of sensitivity. This changed from our preregistered plan to calculate the weighted composite scores, due to recommendations from Strauss et al. (2023)^38^ and Whiteford et al. (2023)^39^ to avoid conflating sensitivity with response bias. All correlations with musical ability were exploratory analyses that predicted a positive relationship between sound neural encoding and melody discrimination and therefore used one-tailed tests. The criterion for significance was Bonferroni-corrected for four comparisons (α = .0125). The /da/ stimulus-to-response correlations used in these analyses used the fixed lag window to match Parbery-Clark et al. (2009)^10^.

### Exclusion criteria

Only those who met the criteria listed in the “Recruitment eligibility” and “Participants” sections took part in the study. Missing data from one or more tasks (e.g., from dropping out of the study or researcher error) resulted in exclusion on the corresponding analyses; whenever possible, the participant was rerun on tests with missing data. Each site has a log of explanations for missing data and technical issues.

EEG data were excluded if there were less than 60% usable trials for any reason, such as a reduced number of trials due to technical issues, researcher error, or an excessive number of artifacts. If a participant did not have enough usable trials, they were re-run on the corresponding task whenever possible. EEG data from the /mi3/ test was excluded if the SNR was too poor to estimate F0 tracking in the sliding FFT analysis. This occurred if the spectral magnitude of the EEG response for all frequencies was within the noise floor.

## Acknowledgments

A.J.O., H.M.B., I.S.J., G.K. Jr., A.E.L., R.K.M., E.W.M., T.K.P., and B.G.S. disclose support for the research of this work from the National Science Foundation [NSF-BCS grant 1840818]. A.J.O. discloses support for the research of this work from the National Institutes of Health [R01 DC005216]. I.S.J and trainees V.I., B.M. and S.vH. were funded by the Canada First Research Excellence Fund Award “BrainsCAN” (2017-2023) to Western University. We thank numerous undergraduate researchers who assisted in data collection and/or data quality management, including Penelope Corbett, Angela Sim, and Kara Stevens.

