## Supplementary Information for "Musical training does not enhance neural sound encoding at early stages of the auditory system: A large-scale multisite investigation"

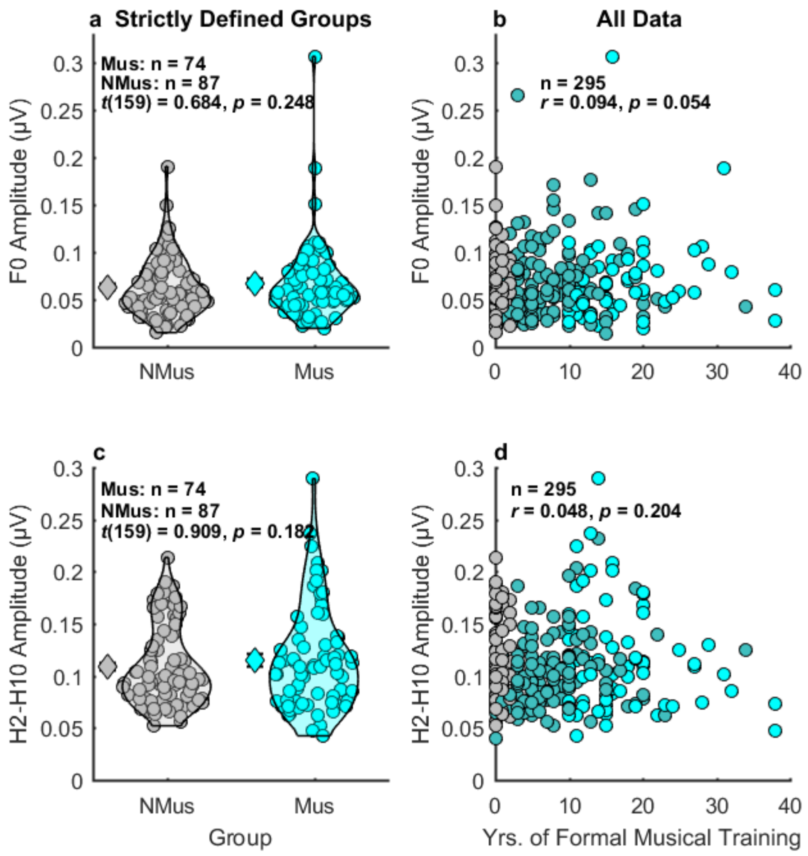


**Supplementary Figure 1. Spectral encoding for /da/ in babble with one non-musician outlier excluded.** There was no musician advantage for encoding the F0 (a,b,) or upper harmonics (c,d). Black outlines: 1D kernel density estimates (KDEs); Diamonds: Average data; Circles: individual data; NMus: non-musicians (grey); Mus: musicians (cyan). Participants in neither of the strictly defined groups are shown in dark cyan.

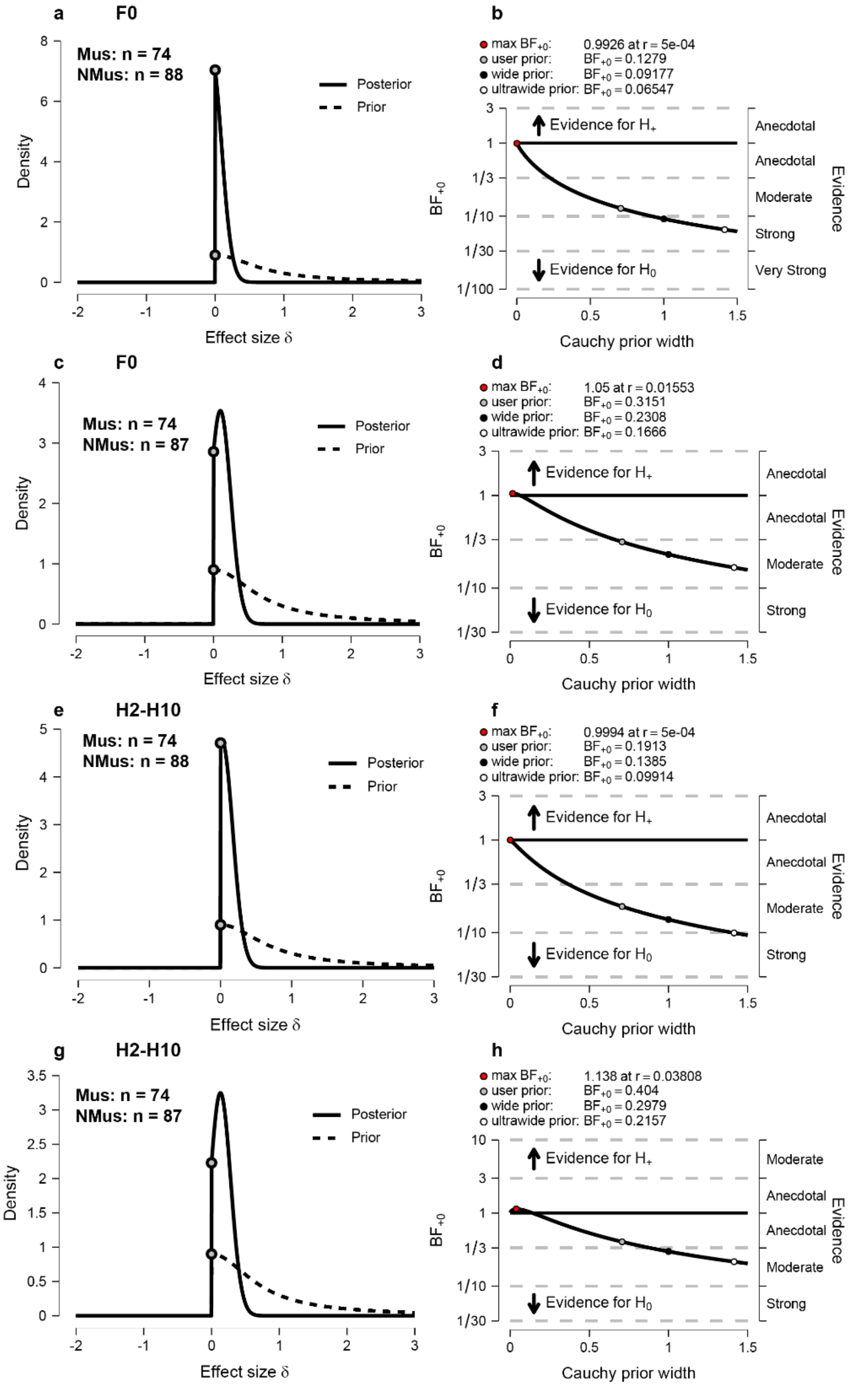


**Supplementary Figure 2. Prior and posterior distributions (a,c,e,g) and robustness checks (b,d,f,h) for Bayesian independent samples *t*-tests comparing spectral encoding for /da/ in babble in musicians vs. non-musicians.** Panels a-d correspond to F0 amplitude and panels e-h correspond to the amplitude of the upper harmonics. Analyses for all musicians and non-musicians (a,b,e,f) versus the same analyses with the best non-musician excluded (c,d,g,h). Robustness checks show the evidence for the null hypothesis is moderate and consistent across reasonable prior widths (*r*). Grey circles: $BF_{+0}$ when $r = \frac{1}{\sqrt{2}}$; black circles: $BF_{+0}$ when $r = 1$; white circles: $BF_{+0}$ when $r = \sqrt{2}$ (b,d,f,h).

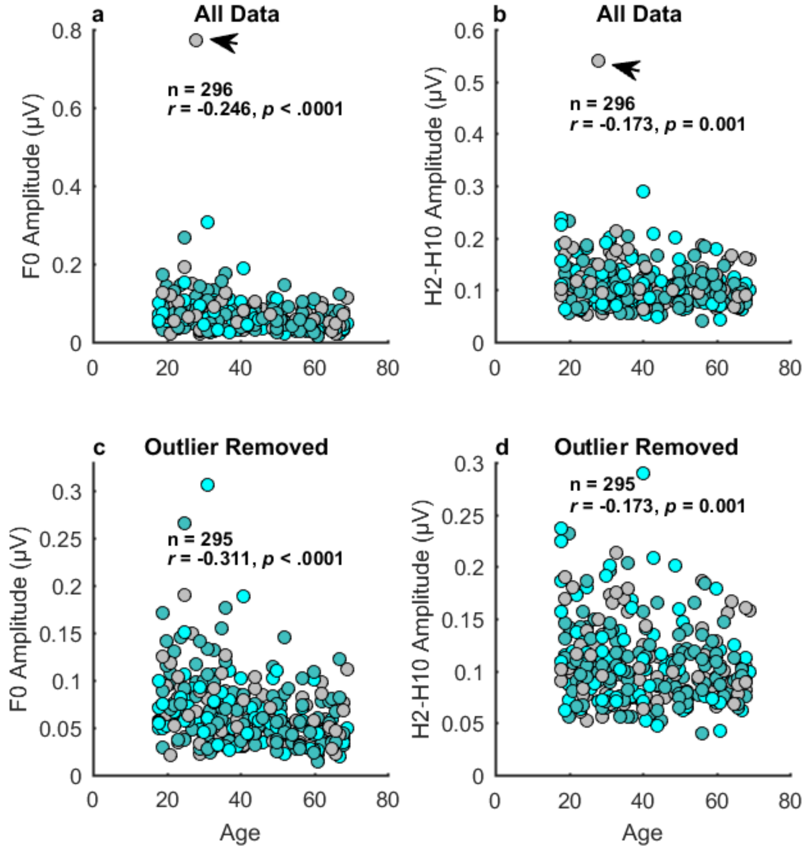


**Supplementary Figure 3. Spectral encoding to /da/ in babble degrades with age.** All data (a,b) and non-musician outlier excluded (c,d). NMus: non-musicians (grey); Mus: musicians (cyan). Participants in neither of the strictly defined groups are shown in dark cyan. Arrow denotes outlier non-musician.

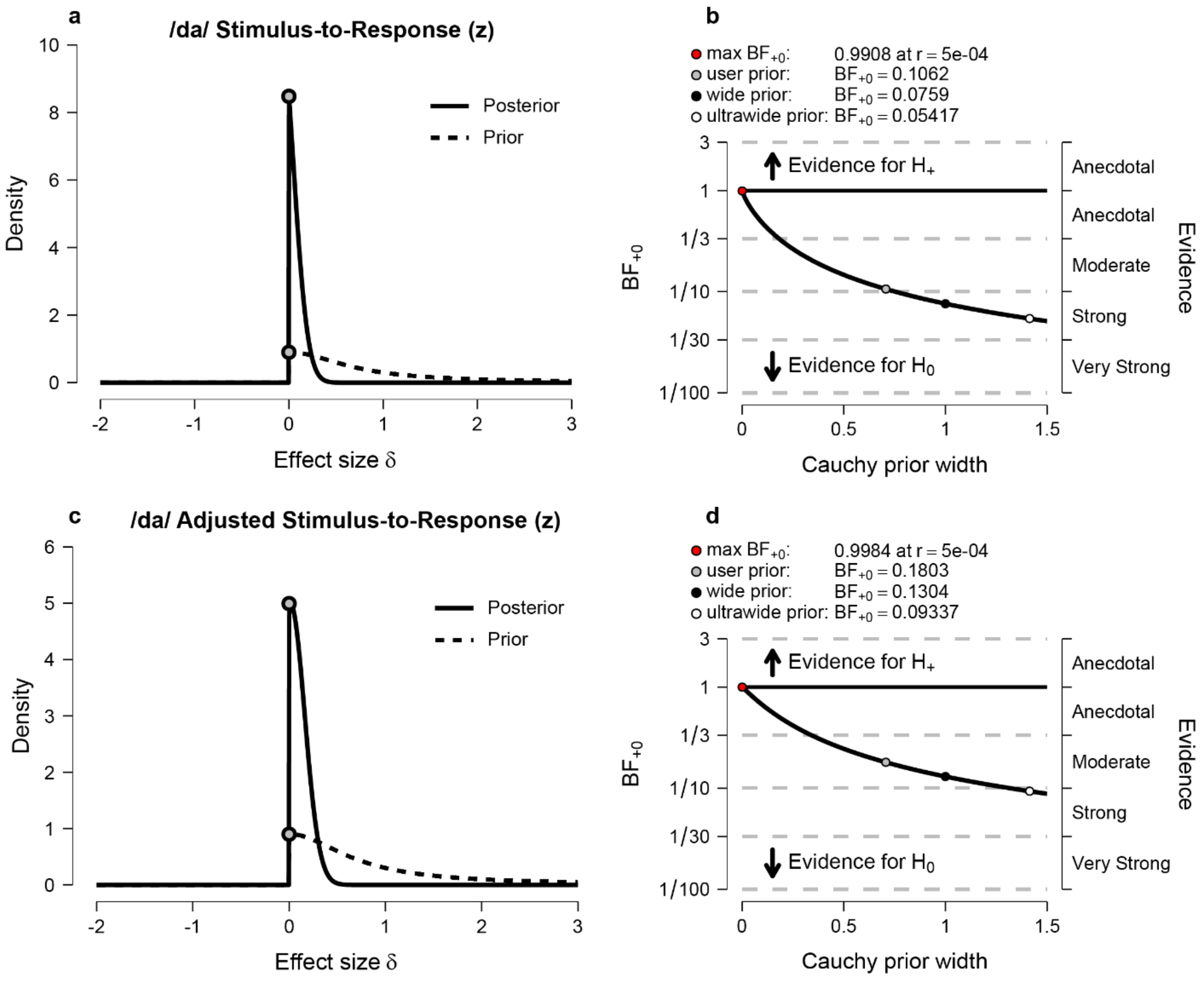


**Supplementary Figure 4. Bayesian independent-samples *t* tests (a,c) and the Bayes factor robustness plots (b,d) for testing whether the fidelity of neural encoding for /da/ in babble is greater in musicians (n=74) relative to non-musicians (n=88).** Neural encoding was quantified as the z-transformed stimulus-to-response correlation for each participant using the preregistered method (a,b) and an alternative method that directly matched the analysis from Parbery-Clark et al. (2009)[1], which limited the cross-correlations to the site-specific adjusted lag times of 6.9-10.0 ms (c,d). Analyses showed strong (a,b) and moderate (c,d) support for the null hypothesis, and results were generally robust to prior width (*r*) . In the robustness plots, the grey circles correspond to the $BF_{+0}$ with the prior selected for analyses ($r = \frac{1}{\sqrt{2}}$), black circles correspond to a wide prior ($r = 1$), and white circles correspond to an ultra-wide prior ($r = \sqrt{2}$); red circles correspond to the prior width with maximal $BF_{+0}$.

### Exploratory Analyses Correlating Musical Training with Neural Encoding for /da/ while Controlling for Age

There was no relationship between years of formal musical training and spectral encoding for the F0 (all participants: $r_p$ = .009, *p* = .439; outlier excluded: $r_p$ = .085, *p* = .072) or upper harmonics of /da/ in babble (all participants: $r_p$ = .002, *p* = .488; outlier excluded: $r_p$ = .042, *p* = .236), nor was there a relationship between musical training and the stimulus-to-response correlation of the vowel portion of the neural response with or without limiting the cross-correlations to the site-specific adjusted lag windows of 6.9-10.9 ms (no adjustment: $r_p$ = -.009, $p$ = .563; site-specific adjustment: $r_p$ = .033, $p$ = .285).

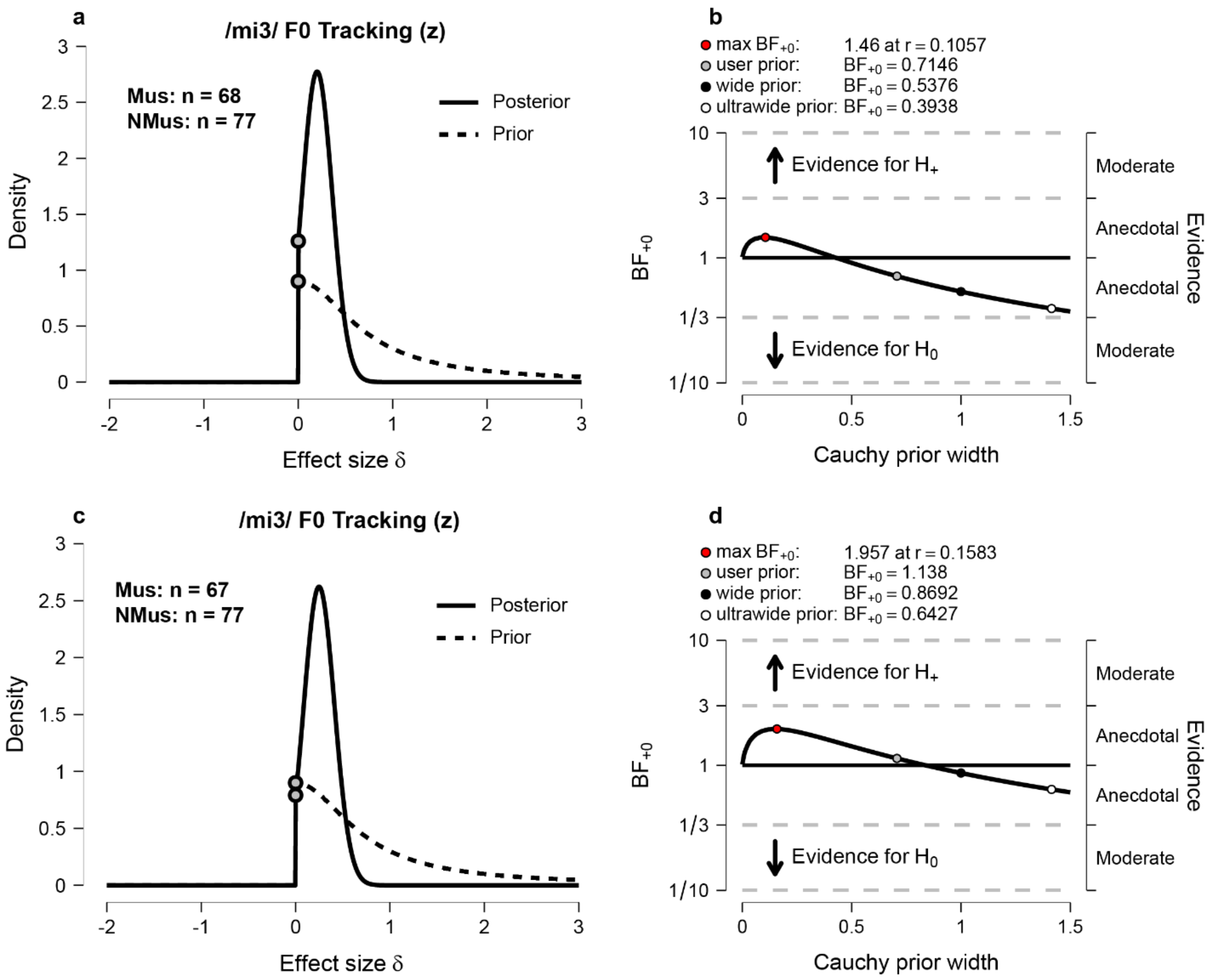


**Supplementary Figure 5. Bayesian independent samples *t* tests (a,c) and robustness checks (b,d) testing whether musicians have stronger F0-tracking for /mi3/ relative to non-musicians.** Evidence slightly favored the null hypothesis before (a,b) but not after removing one outlier musicians (c,d). Overall, there was no strong evidence for either the alternative or the null hypothesis.

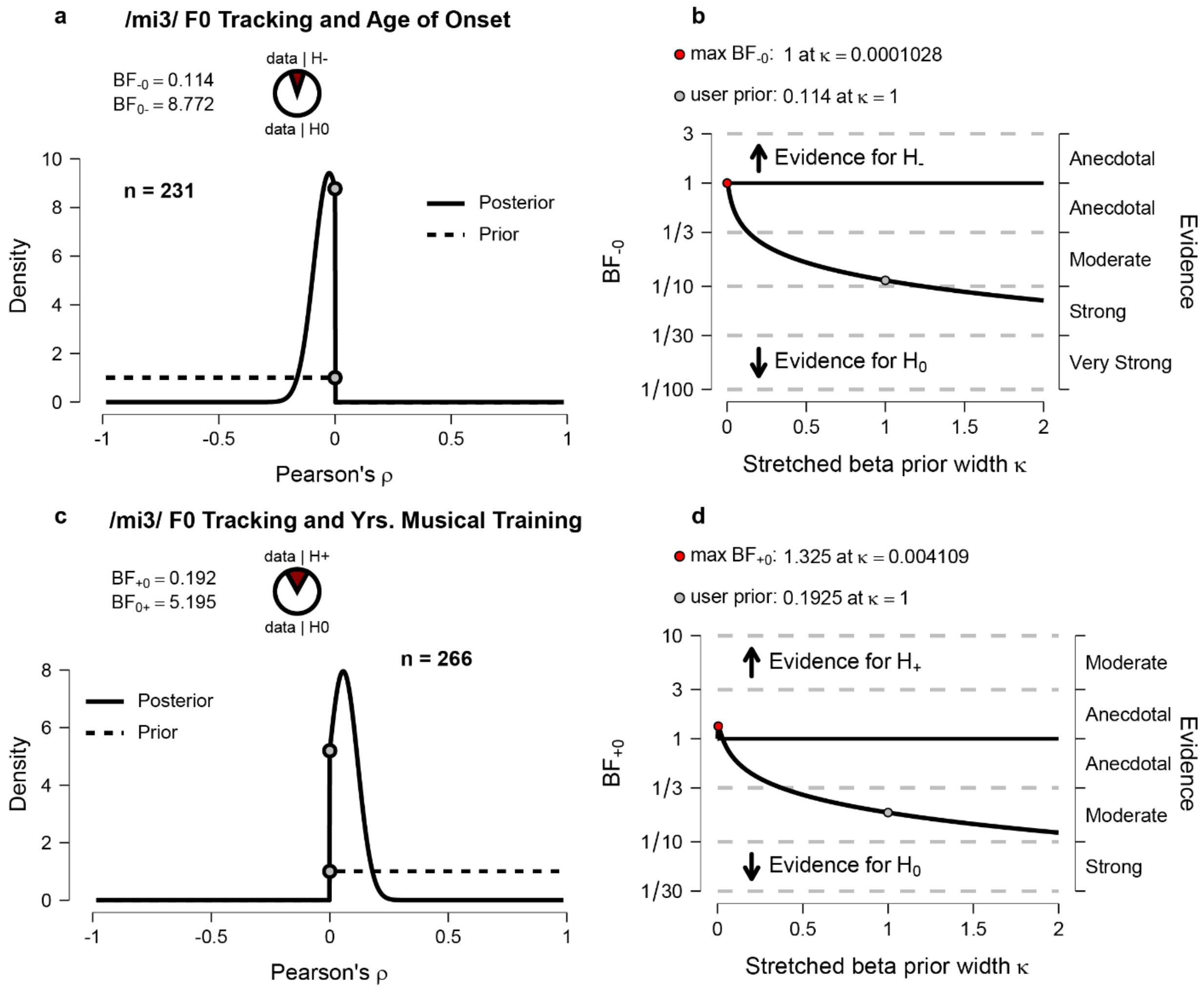


**Supplementary Figure 6. Bayesian Pearson correlations (a,c) and Bayes factor robustness plots (b,d).** Tests the hypothesis that the strength of F0-tracking is greater the earlier one started playing their first musical instrument or voice (a) and the hypothesis that F0-tracking increased with increased years of formal musical training (c). In both instances, there was moderate evidence in support of the null hypothesis, and this interpretation was generally robust to varying prior widths for both age of onset (b) and years of musical training (d).

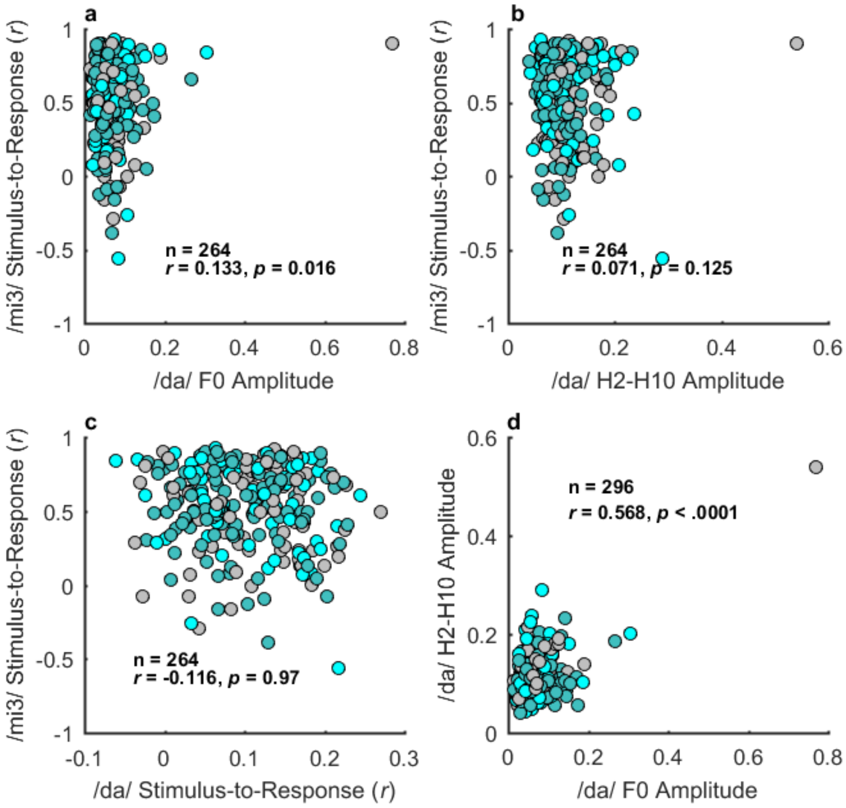


**Supplementary Figure 7. Pairwise correlations of four FFR measures: All participants.** One outlier non-musician had unusually strong spectral encoding.

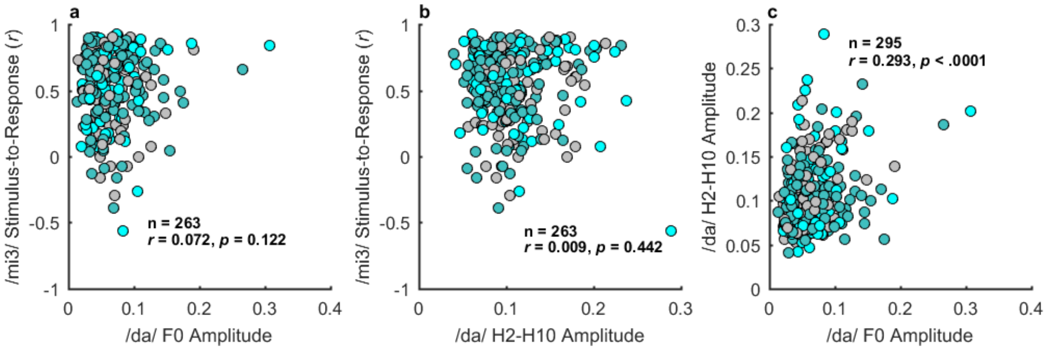


**Supplementary Figure 8. Pairwise correlations of four FFR measures: Outlier non-musician excluded.**

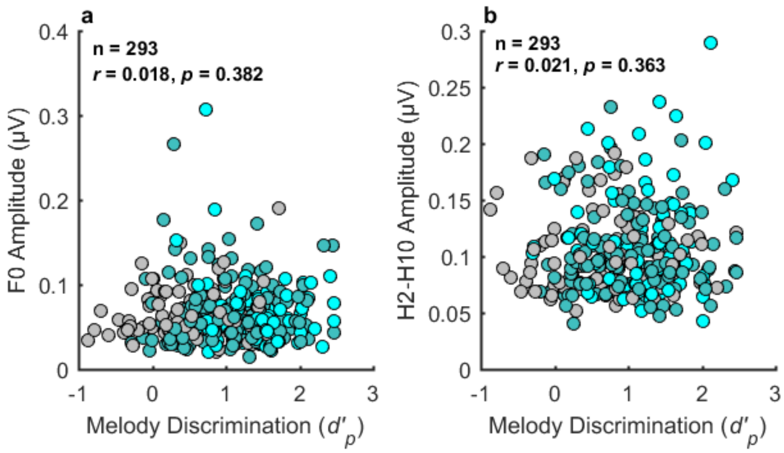


**Supplementary Figure 9. No relationship between musical ability and spectral encoding for /da/ in babble for the F0 (a) or upper harmonics (b), with one non-musician outlier removed.**

**Supplementary Table 1. Number of participants per site that were included in analyses.**

**A. /da/ in Babble**

| | **Site** | | | | | |
|---|---|---|---|---|---|---|
| **Group** | **BU** | **CMU** | **PU** | **UMN** | **UR** | **UWO** |
| Mus. | 9 | 5 | 6 | 29 | 16 | 9 |
| NMus. | 17 | 14 | 10 | 18 | 12 | 17 |
| Variable Mus. | 12 | 14 | 10 | 42 | 35 | 21 |

**B. /mi3/ in Quiet**

| | **Site** | | | | | |
|---|---|---|---|---|---|---|
| **Group** | **BU** | **CMU** | **PU** | **UMN** | **UR** | **UWO** |
| Mus. | 9 | 5 | 5 | 26 | 15 | 8 |
| NMus. | 15 | 11 | 8 | 16 | 11 | 16 |
| Variable Mus. | 10 | 14 | 8 | 40 | 30 | 19 |

*Note.* Non-Mus. = Non-Musician; Mus. = Musician; Variable Mus. = Variable Amounts of Musical Training.

### Post-Registration Changes to Methods

#### General notes

One site (PU) did not maintain EEG logs detailing any deviations from the protocol that occurred. Any notes present were compiled by KLW by communicating directly with the site.

#### Melody discrimination

1. The preregistration incorrectly stated that we would use the Melody subtest from the Mini Profile of Music Perception Skills (Mini-PROMS[2]); the Melody trials used during piloting and the full study were from the Full PROMS[3].

2. Melody discrimination performance was analyzed using a non-parametric estimate of sensitivity, $d'_p$, as recommended by Strauss et al. (2023)[4] and Whiteford et al. (2023)[5] to avoid issues of conflating sensitivity with response bias that occur with the weighted composite score.

3. Two participants from UR (s45 and s58) indicated they had audio issues on the melody discrimination task but were not excluded from the experiment and were not rerun on the melody task in the lab. Data from these participants on the other measures from our study are still included, except for analyses involving the melody discrimination measure, for which they were excluded.

**EEG**

Five subjects (s18, s19, s20, s21, and s23) from CMU were run with left and right mastoid as the references due to experimenter error, rather than the left and right earlobes. These subjects were included in all analyses.

**Participants**

Each site had a preregistered recruitment goal of n=60 participants, consisting of 15 musicians, 15 non-musicians, and 30 that did not fit either group, with participant age spread with a uniform distribution. Deviations from the initial recruitment goals occurred due to practical constraints, including the COVID-19 pandemic.

### Post-Registration Changes to Analysis Plan

1. Author KLW assisted with cleaning and quality-checking data from site PU.

2. Author PYG assisted author KLW with preprocessing EEG data from all sites.

3. Author KLW wrote the code for cleaning and formatting the raw EEG data for each site. One exception was the initial formatting of the UR data to a .mat file, which was done by the staff at that site location. Each site was still responsible for ensuring their EEG data were suitable for inclusion in the analyses.

4. Most of the participants in the present study also participated in multiple preregistered behavioral experiments related to musical training and sound perception. The behavioral analyses, along with analyses linking behavior and neural coding, will be published in a separate manuscript focusing on the relationship between musical training and sound perception.

5. The pilot data had the incorrect fixed delay (i.e., the delay between the onset of the trigger and the arrival time of the stimulus at the ear canal) for site PU. The analysis code uses the correct fixed delay (1.9 ms).

6. Pre-planned multi-channel analyses were not conducted due to a recent publication demonstrating that complex principal components analysis of the phase-locking values (PLV) across all electrodes is equivalent to taking the average of the squared PLV[6].

**EEG responses to /da/ syllable in noise:** Parbery-Clark et al. (2009)[1]

***Encoding of upper harmonics using FFT.*** After calculating the average spectral amplitudes within each bin separately for H2-H10, the summed amplitude was taken across these averaged bins, rather than the grand average. This decision was made to exactly match the analysis from the original study. The preregistration initially planned to analyze the data using a 2 x 2 repeated-measures ANOVA. However, the use of the summed amplitude for the upper harmonics makes the comparison between the two conditions (F0 vs. upper harmonics) less comparable. Therefore, we chose to use two separate t-tests to analyze the effects of spectral encoding on musical training, which was identical to the analyses used by the original study.
